# The Plateau Method for Forensic DNA SNP Mixture Deconvolution

**DOI:** 10.1101/225805

**Authors:** Darrell O. Ricke, Joe Isaacson, James Watkins, Philip Fremont-Smith, Tara Boettcher, Martha Petrovick, Edward Wack, Eric Schwoebel

## Abstract

Identification of individuals in complex DNA mixtures remains a challenge for forensic analysts. Recent advances in high throughput sequencing (HTS) are enabling analysis of DNA mixtures with expanded panels of Short Tandem Repeats (STRs) and/or Single Nucleotide Polymorphisms (SNPs). We present the plateau method for direct SNP DNA mixture deconvolution into sub-profiles based on differences in contributors’ DNA concentrations in the mixtures in the absence of matching reference profiles. The Plateau method can detect profiles of individuals whose contribution is as low as 1/200 in a DNA mixture (patent pending)^1^.

## Introduction

DNA analysis is a common tool used within law enforcement to identify contributors to forensic evidence. However, crime scene evidence often contains DNA from multiple individuals, which confounds current DNA analysis techniques. Currently, the forensics community identifies individual DNA samples through analysis of short tandem repeats (STRs) which are processed using capillary electrophoresis. This methodology has been proven accurate for identification searches of an individual sample versus a reference database. STR analysis of DNA mixtures with reference samples works for samples with two contributors where the ratio of DNA concentrations does not exceed 1:10, except in the case of rape samples where special techniques are applied to isolate the perpetrator’s cells^2^. PCR amplification of STR repeat structures introduces stutter peak artifacts^3^ that confound the discrimination of contributor STR alleles. Multiple mixture analysis and deconvolution methods have been developed for STR mixture analysis^4^ including: linear mixture analysis method (LMA)^5^, least-square deconvolution^6^, binary models, and semi-continuous and continuous probabilistic methods^7^. Available STR solutions include: ArmedXpert^8^, DNAmixtures^9^, DNA VIEW Mixture Solution^10^, GeneMarker HID^11^, FST^12^, GenoProof Mixture^13^, GenoStat^14^, Lab Retriever^15^, LikeLTD^16^, LiRa^17^, LiRaHt, LRmix Studio^18^, MixSep^19^, STRmix^20^, and TrueAllele^21^. While multiple methods claim to handle mixtures of more than two individuals, the state of the art continues to be that multiple experts develop different interpretations of mixture data^22^.

One possible solution to the limitation imposed by the stutter problem is to replace or augment STRs with thousands of SNPs for the purpose of mixture analysis and deconvolution^23^,^24^. However, since there are no current analysis methods for mixture deconvolution using SNPs, we introduce the Plateau method for those mixtures with modest numbers of contributors and imbalanced contributor DNA concentrations.

## Method

### Ion Torrent HTS Sequencing

Swabs (Bode cat# P13D04) were used to collect buccal cells from the inside of cheeks of volunteers, rubbing up and down for at least 10 seconds, with pressure similar to that used while brushing teeth. DNA was isolated from swabs using the QIAamp DNA Investigator Kit (QIAGEN cat#56504), using the “Isolation of Total DNA from Surface and Buccal Swabs” protocol, and eluted in 100uL of low TE (carrier RNA not used; low TE has 0.1mM of EDTA). Quantitation was done using Quantifilier HP kit (Thermo Fisher Scientific cat#4482911) according to manufacturer, with the exception that human genomic DNA from Aviva Systems Biology (cat#AVAHG0001) was used for the standard. Purified DNA was combined to produce defined mixtures. Primers for 2,655 targets were designed using the Ion AmpliSeq Designer online tool. Libraries were prepared using the AmpliSeq 2.0 library kit protocol according to the manufacturer, with the exception that 19 cycles were performed (no secondary amplification) and the library was eluted in 25uL low TE. Library quantitation was performed using the Ion Library Quantitation Kit (Thermo Fisher cat#4468802), according to the manufacturer. Template preparation and sequencing were performed using the Ion Chef and Ion Proton according to the manufacturer (Thermo Fisher Ion Chef and Proton cat#A27198 and Proton chips cat#A26771).

### HTS SNP Data Analysis

The GrigoraSNPs^25^ program was used to call SNP alleles from multiplexed HTS FASTQ sequences. Mixture analysis was performed using the MIT Lincoln Laboratory IdPrism HTS DNA Forensics system.

### Plateau Method

The Plateau method is based on the observation that the percent contribution of an individual’s DNA in a mixture is reflected by the minor allele ratio (mAR - the ratio of observed minor alleles divided by the sum of the minor and major alleles for each individual SNP), which is calculated for each locus. The loci are sorted by ascending log(mAR) and sub-profiles are identified by an approach that identifies clusters of SNPS in plateaus with very similar minor allele ratios. The DBSCAN^26^ algorithm is used to cluster the loci using the log difference in mAR between adjacent loci (sorted by mAR) as the distance metric. For a panel of 2655 loci, a minimum cluster size of 10 and an epsilon of 0.001 achieve reliable separation between a mixture’s contributors. The DBSCAN algorithm identifies loci that cannot be assigned to any clusters in the data. These loci are used to demarcate the beginning and end of a single individual’s discernable major:minor alleles contribution to a mixture. SNPs below a minimum mAR threshold are classified as homozygous major for all identified sub-profiles. When multiple sub-profiles are identified, the minor allele loci of sub-profiles to the left of a given sub-profile, i.e. those with lower average mAR, are classified as homozygous major for that sub-profile. Loci with mAR greater than 0.99 in the mixture are assigned to the sub-profile with the highest mAR cluster.

### Results and Discussion

A reference profile consists of homozygous major (MM), heterozygous (mM), and homozygous minor (mm) genotypes, where the minor allele ratio (mAR) values at or near 0 represent MM genotypes, near 0.5 represent mM genotypes, and 1.0 represent mm genotypes (Figure 1). The sorted log(mAR) plots for two reference profiles used in the two-person dilution experiments are illustrated in Figure 1. The identified plateaus for the two-person mixture dilution series (DNA ratios 25:75, 10:90, 1:99, 1:200, and 1:400) are shown in Figure 2. The loci with mM genotypes create plateaus of similar mAR values. These plateaus are leveraged by the Plateau method to create individual DNA signatures, or sub-profiles, directly from the mixture without use of reference profiles, resulting in direct deconvolution of sub-profiles from the mixture.

**Figure 1.**
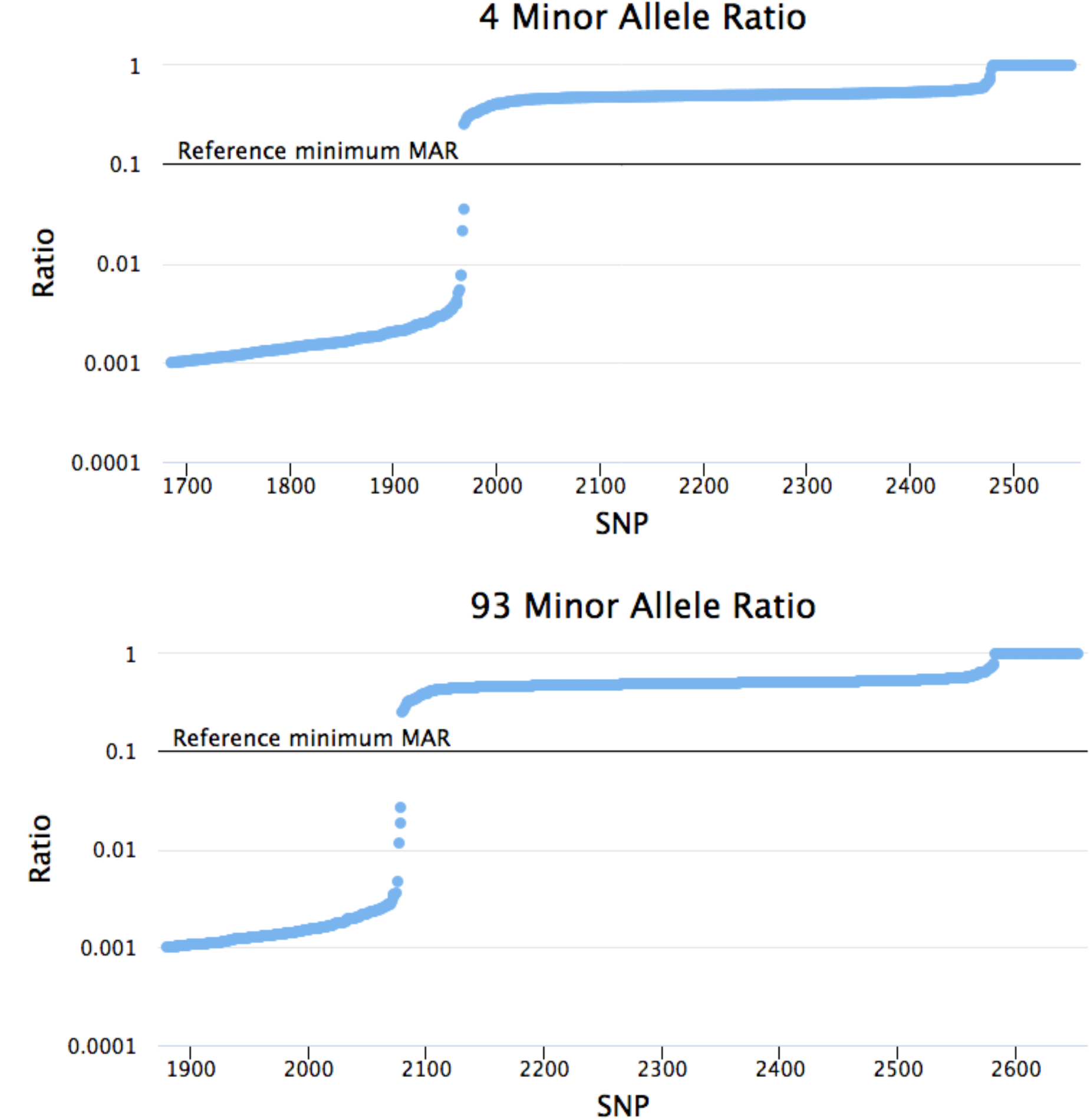
Sorted minor allele ratios for reference profiles for individuals 4 and 93. Profiles are sorted by ascending log(mAR) – MITLL IdPrism screenshots.

**Figure 2.**
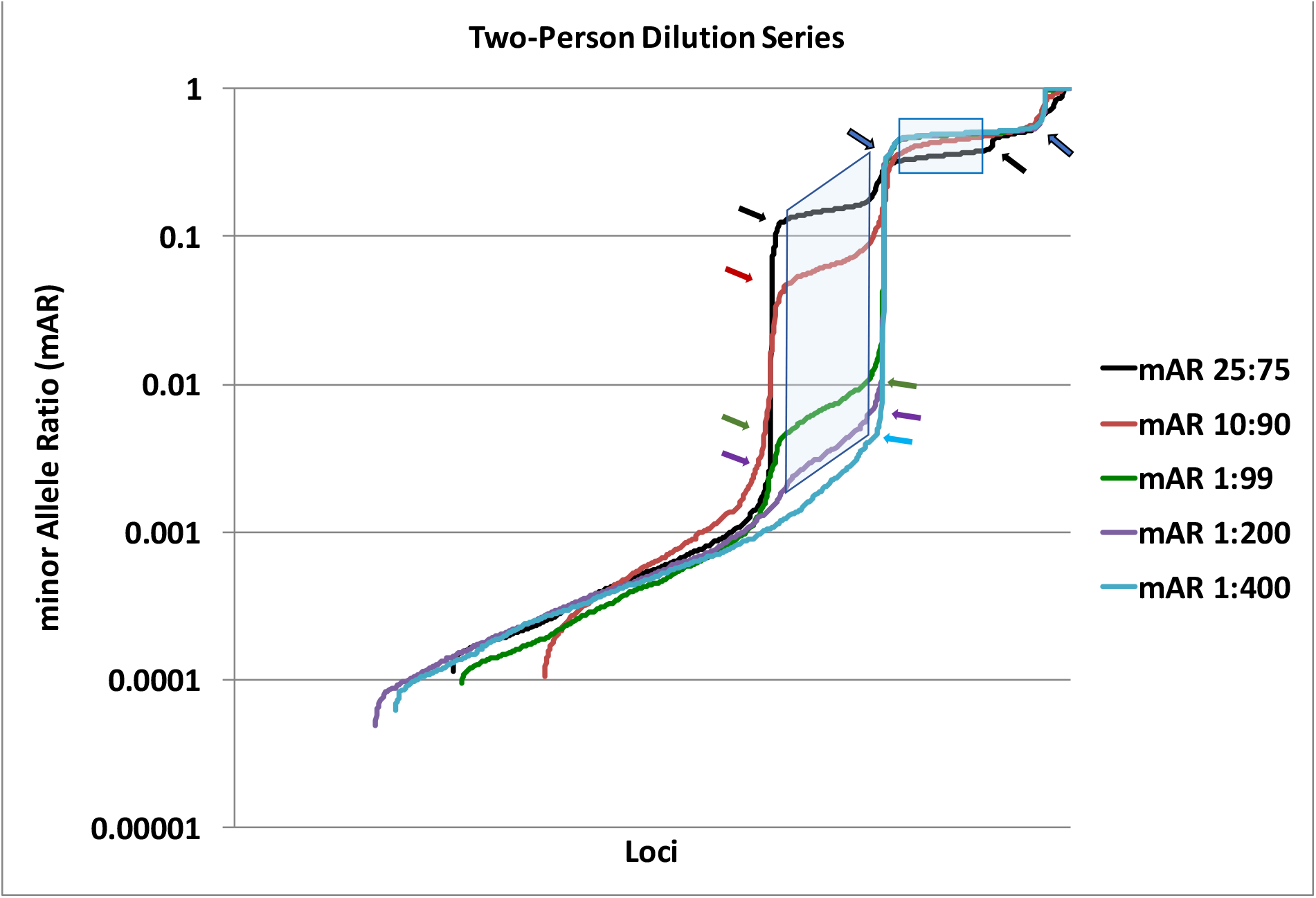
Sorted mAR two-person dilution series of individuals 4 and 93. The boxes highlight the mM allele plateaus in each mixture. Colored arrows point to locations of slope changes that are the outer boundaries for plateau detection; arrow colors correspond to line colors and blue arrows with black outlines are used to indicate multiple overlapping arrow positions. Note that that the lowest light blue arrow for 1:400 is missing as there is a lack of significant change in mAR line slope.

Individual profiles can be resolved with DNA ratios as low as 1:200 for both DNA contributors. For the 1:400 mixture, a plateau is detected for the major DNA contributor, but the shape of the mM alleles for the lower DNA contributor does not qualify cleanly as a plateau because there is a lack of significant slope change between the lowest contributor’s mAR values and loci with putative noise values. Figures 3 and 4 illustrate the Plateau method applied to defined mixtures of three and four individuals, respectively. Table 1 compares the individual sub-profiles identified by the Plateau method to the truth data the two-person dilution series, three-person dilution mixture in Figure 3, and four-person dilution mixture in Figure 4. The Plateau method can identify plateaus for some mixtures with more than two contributors as long as there is a sufficient difference in contributor DNA concentrations (≥ 15%), (see Figures 3 & 4 with the difference between each individual contributor is roughly 15%). The Plateau method creates highly enriched sub-profiles that uniquely identify contributors with low probability of random man not excluded (P(RMNE)) values even when rare combinations of minor SNP alleles from other DNA contributors are included in the plateaus (See Table 1, 4-person mixture, individual 93).

**Figure 3.**
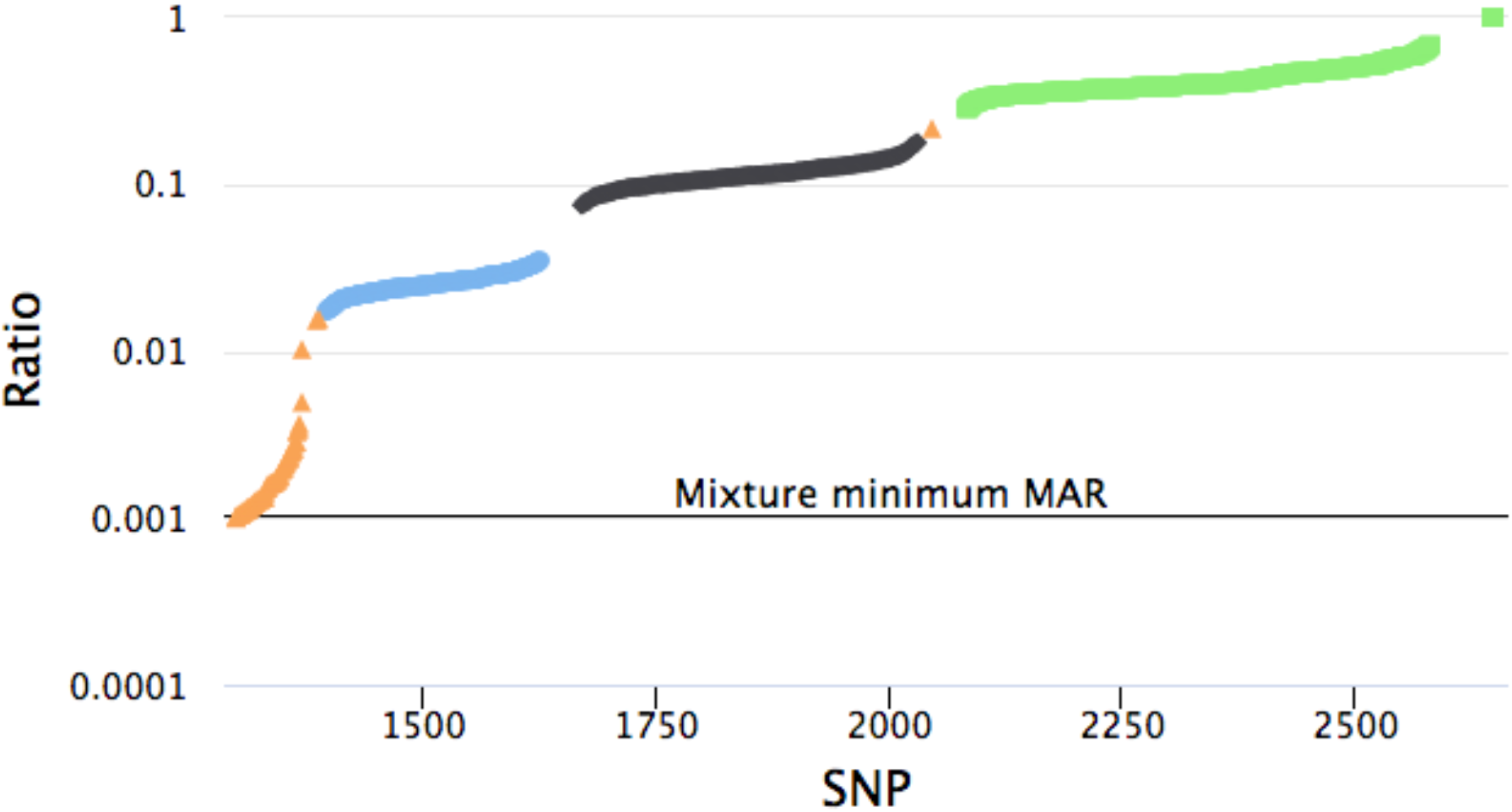
Three-person mixture with DNA concentrations individual 56 at 5%, individual 24 at 20%, and individual 93 at 75% (MITLL IdPrism screenshot)

**Figure 4.**
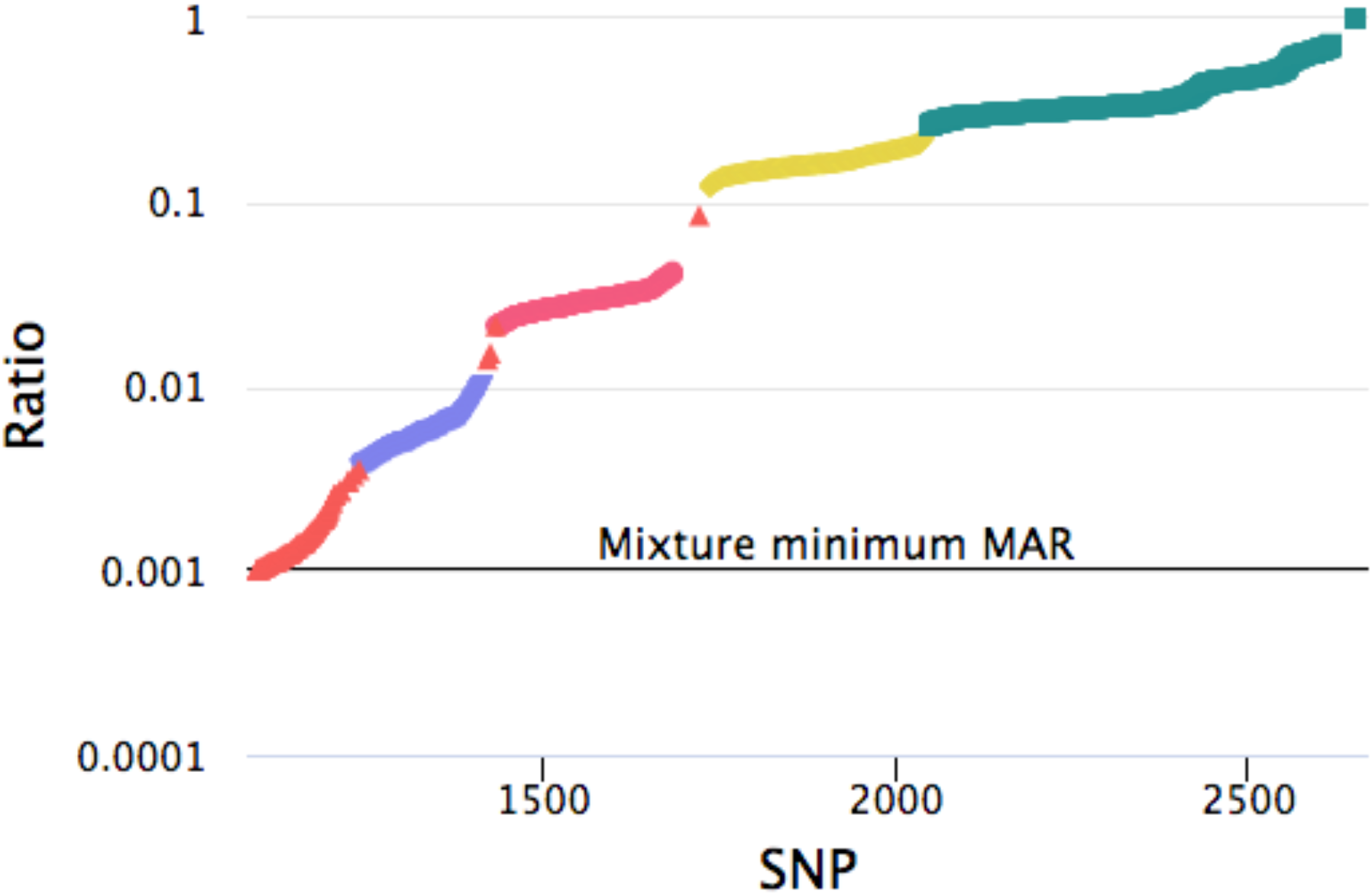
Four-person mixture with DNA concentrations individual 4 at 2%, individual 94 at 17%, individual 78 at 32%, and individual 93 at 47% (MITLL IdPrism screenshot)

**Table 1.** Analysis of Plateau sub-profiles versus Reference Samples.

| Mixture | Number of Contributors | Individual | MM alleles | mM alleles | mm alleles | mismatches | P(RMNE) |
| --- | --- | --- | --- | --- | --- | --- | --- |
| 25:75 | 2 | 4 | 2018 | 314 | 35 | 26 | 5.92E-107 |
| 25:75 | 2 | 93 | 1727 | 256 | 41 | 2 | 8.25E-103 |
| 10:90 | 2 | 4 | 1986 | 461 | 27 | 1 | 4.72E-138 |
| 10:90 | 2 | 93 | 1721 | 238 | 39 | 6 | 5.33E-106 |
| 1:99 | 2 | 4 | 1954 | 430 | 27 | 0 | 1.87E-143 |
| 1:99 | 2 | 93 | 1670 | 257 | 39 | 0 | 9.03E-114 |
| 1:200 | 2 | 4 | 1855 | 447 | 34 | 0 | 6.82E-134 |
| 1:200 | 2 | 93 | 1631 | 190 | 45 | 5 | 1.90E-100 |
| 1:400 | 2 | 4 | 1901 | 477 | 0 | 0 | 5.20E-144 |
| 1:400 | 2 | 93 | 1660 | 241 | 0 | - | - |
| 5:20:75 | 3 | 93 | 1959 | 474 | 24 | 0 | 2.02E-142 |
| 5:20:75 | 3 | 56 | 1359 | 222 | 0 | 2 | 1.95E-78 |
| 5:20:75 | 3 | 24 | 1581 | 355 | 0 | 1 | 1.85E-102 |
| 2:17:32:47 | 4 | 4 | 1182 | 175 | 0 | 2 | 3.81E-61 |
| 2:17:32:47 | 4 | 93 | 1935 | 549 | 22 | 42 | 8.73E-91 |
| 2:17:32:47 | 4 | 94 | 1357 | 251 | 0 | 0 | 5.65E-84 |
| 2:17:32:47 | 4 | 78 | 1608 | 306 | 0 | 7 | 7.00E-90 |

When tested on four- and five-person mixtures with differences of only 10% in DNA concentration, the Plateau method only identifies a single plateau that corresponds with the 10% DNA contributor (see Figure 5). For these three mixtures, the slope of the mAR of the three contributors with higher than 10% DNA concentration do not readily demarcate. In addition, the profile for the 0.5% DNA contributor in the 5-person mixture is visible in Figure 5, but is not classified as a plateau due to the high slope angle (blue line between mAR 0.001 and 0.01).

**Figure 5.**
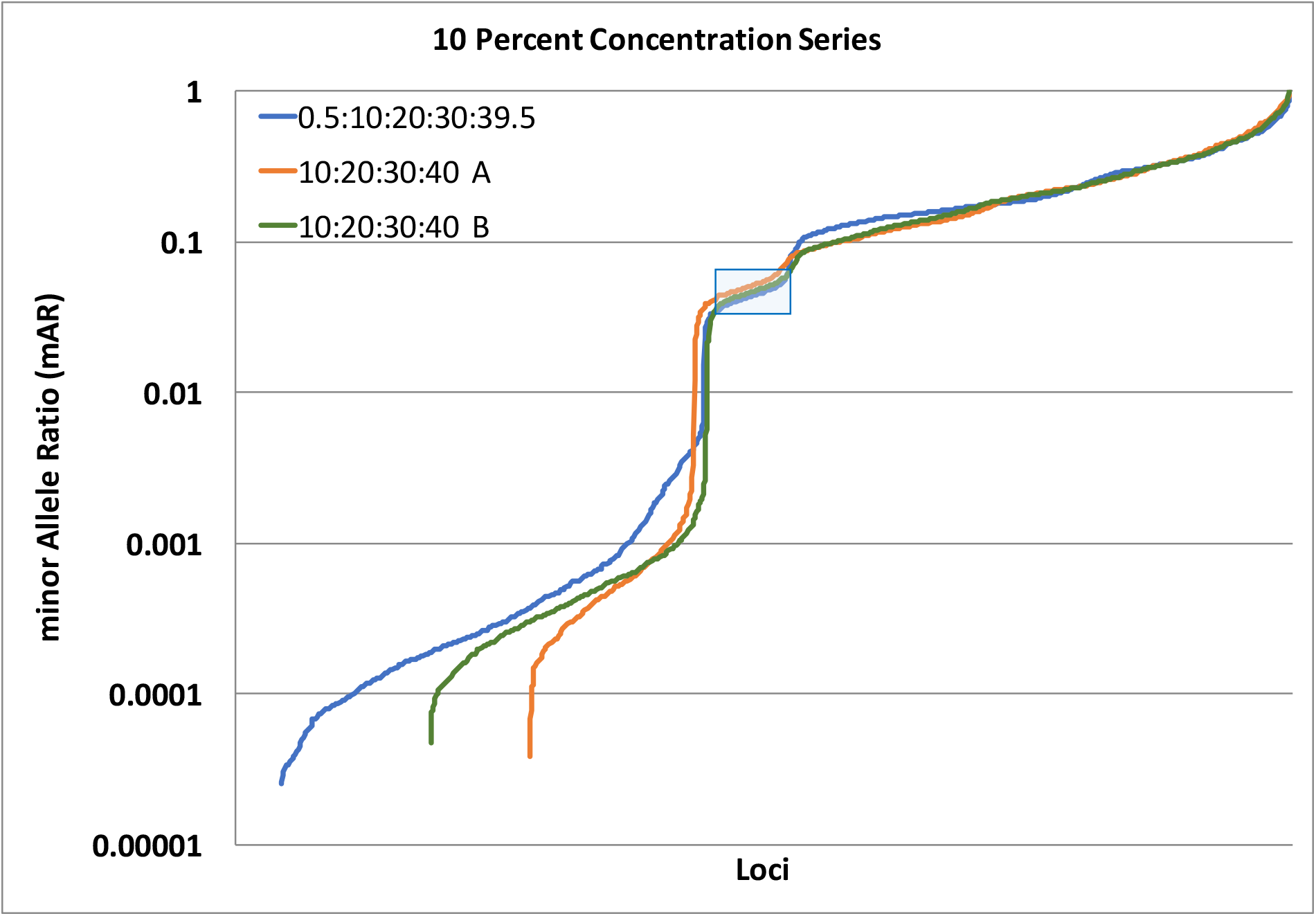
Sorted 10% concentration dilution series for four and five people. The two four-person mixtures use DNA concentrations of 10%, 20%, 30%, and 40%. The five-person mixture uses DNA concentrations of 0.5%, 10%, 20%, 30%, and 39.5%. The Plateau method identifies one sub-profile (boxed) in each of these mixtures that corresponds with the 10% DNA concentration contributor.

The Plateau method enables the direct deconvolution for some mixtures into sub-profiles based on differences in individual mARs within the mixture. This allows the sub-profiles to be matched to other forensic samples, even when no reference profiles exist, thereby enabling linking different events or crime scenes together. The results presented illustrate that the Plateau method can differentiate individuals with ≥ 15% DNA concentration difference and identify individuals down to contributions as low as 0.5% (1:200) contribution, and can resolve mixtures of 2, 3, and 4 individuals. The Plateau method provides a solution for direct deconvolution of some DNA mixtures into individual sub-profiles.

### Conclusion

The Plateau method can directly deconvolve some SNP DNA mixture into sub-profiles for individual contributors without the use of reference profiles. This is the first SNP mixture deconvolution method to be described and compliments DNA mixture analysis methods that match known references to DNA mixtures.

## Acknowledgements

Distribution statement: pending

This material is based upon work supported under Air Force Contract No. FA8721-05-C-0002 and/or FA8702-15-D-0001. Any opinions, findings, conclusions or recommendations expressed in this material are those of the author(s) and do not necessarily reflect the views of the U.S. Air Force.

